# Measuring and removing near-duplicate contamination in alignment-free SARS-CoV-2 lineage classification benchmarks

**DOI:** 10.64898/2026.08.12.744560

**Authors:** Mohammad Jamhuri, Andy Irawan

## Abstract

Alignment-free lineage assignment from *k*-mer frequency profiles is widely used for SARS-CoV-2 surveillance, and the methods that do it are ranked against each other by margins of one or two percentage points. Those rankings rest on an unchecked protocol. Public repositories hold many near-duplicate genomes, and stratified random splitting puts members of such a group on both sides of the split, so a classifier is credited for sequences it has already seen. We propose quantised profile hashing, which finds near duplicates in *k*-mer feature space by rounding each frequency vector and hashing it. No sequence is compared with any other, so one pass over the feature matrix suffices and no similarity threshold has to be chosen. Rounding is also what makes the groups well defined, and they are then kept whole across the training, validation and test sets. On 255,611 genomes from seven Pango lineages, random splitting leaves 5.09% of test sequences with a near duplicate in training, on a benchmark ranked by margins of one or two points. Ten update rules were trained twice, identically except for the partition. The contaminated benchmark separates one rule from the leader at 0.05; the clean one separates none. The two orderings are uncorrelated, Kendall *τ* = +0.022, with rules moving 3.2 positions on average and the leader of one benchmark ranking eighth on the other. A ranking obtained under contamination therefore says nothing about the ranking without it, and the quantity worth reporting beside a score is the leakage rate of the split.

## 1. Introduction

SARS-CoV-2 is still evolving. Assigning a new genome to its lineage remains a daily task in genomic surveillance, and public repositories now hold millions of genomes. The usual tools compare each genome against reference sequences or lists of marker mutations. They are accurate, but their cost grows with the size of the reference set, and recombination breaks the branching tree they assume.

Alignment-free methods avoid this. Each genome becomes a fixed-length vector describing its composition, usually the frequency of its *k*-mers, with 4^*k*^ entries whatever the genome length and computed in one pass. Randhawa et al. [1] combined supervised learning with digital signal processing; Sung et al. [2] proposed AutoCoV, Cacciabue et al. [3] developed Covidex, and Miao et al. [4] reviewed the field. At larger scale, van Zyl et al. [5] reached 97.8% accuracy on 297,186 sequences over 3,502 lineages, and Yu et al. [6] a macro F1-score of 0.9636 over 103 lineages. Other work encodes genomes as images [7, 8, 9] or applies convolutional models to raw sequences [10].

Two things stand out in this literature. Reported scores lie in a narrow band, so methods are ranked by margins of one or two percentage points; and almost every study divides its data by stratified random splitting. That second habit is a known hazard for biological sequences. Repositories hold many sequences that are nearly identical, and a random split puts such sequences on both sides at once, so a model can score well by recognising what it has already seen. Bernett et al. [11] retrained leading protein–protein interaction models on leakage-free splits and saw their scores fall sharply. Tools for safer splitting have followed, among them SpanSeq [12] and DataSAIL [13]. Both compute pairwise similarity between sequences and then cluster: SpanSeq with *k*-mer sketch distances and DBSCAN at minPoints = 1, which reduces to single-linkage clustering, and DataSAIL with MM-seqs2 or Needleman–Wunsch alignment. DataSAIL also defines a quantitative measure of information leakage.

Two questions follow for SARS-CoV-2 lineage classification, and neither has been answered. How much near-duplicate contamination does a randomly split benchmark contain? And does removing it change what the benchmark reports? The second does not follow from the first: a contamination too small to move any published score could still be enough to re-arrange the order of the methods being compared, and it is the order that such benchmarks are read for.

This paper answers both on 255,611 genomes from seven lineages. We propose *quantised profile hashing* (QPH), which finds near duplicates in *k*-mer feature space: frequency vectors are rounded and hashed, and sequences sharing a digest form one group that is never split. Rounding is what makes the groups well defined — a distance threshold would not, because closeness does not pass along a chain — and no sequence is compared with any other, so one pass over the feature matrix suffices. Measured this way, stratified random splitting leaves 5.09% of test sequences with a near duplicate in training, on a benchmark ranked by margins of one or two points. Ten update rules were then trained twice, on the contaminated split and on the clean one, identically in every other respect. The contaminated benchmark separates one rule from the leader; the clean one separates none, and the two rankings are uncorrelated. Following DataSAIL [13], which quantifies leakage but does not report it for this task, we give *λ* for both partitions and argue that it belongs beside any published score. The code, both splits and the sixty trained models are released, so the check can be repeated elsewhere without retraining.

Three points set this apart from the work it departs from. First, SpanSeq and DataSAIL reach their groups by comparing sequences and clustering the result, whereas QPH does neither; and it compares the feature vector the classifier is trained on rather than a sketch of the sequence, which is the strictest place at which two inputs can be called the same. Second, single-linkage clustering is the transitive closure of a threshold relation, so a chain of intermediate sequences can merge two profiles that are far apart and the outcome depends on the threshold; rounding is transitive by construction and cannot chain. Third, Bernett et al. [11] found that protein interaction models fall to chance once leakage is removed and are then matched by simple baselines. Nothing of the kind happens here: every rule stays above a macro F1-score of 0.995 on both partitions and the scores move by less than 0.002, yet the two rankings are uncorrelated. Leakage can therefore empty a benchmark of information while leaving no visible drop in performance, which is the harder case to detect.

## 2. Methodology

Figure 1 shows the whole procedure in the order it was carried out, and this section follows that order: first the sequences are prepared, then a genome becomes a vector, near duplicates are found and the split is built, and finally the classifier and the ten update rules are described. What the paper proposes lies in the middle part — the QPH criterion of Subsection 2.4, the split constraint built on it, and the leakage rate *λ*. The classifier and the rules are the instrument used to ask whether the split changes what a benchmark reports, not proposals of their own, and are given in full only so that a reader can check that the instrument is sound.

**Figure 1:**
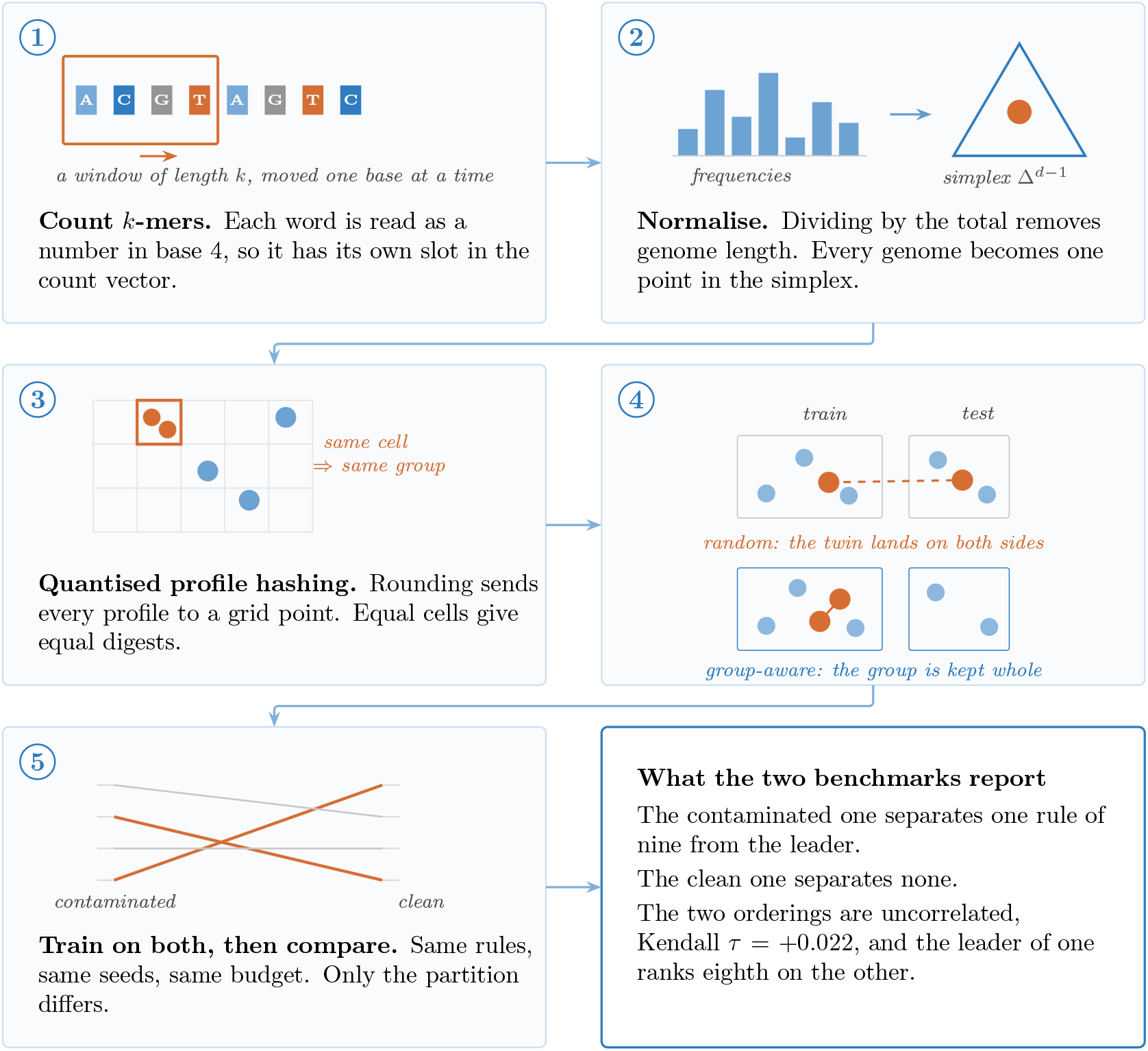
How the study proceeds, read left to right and top to bottom. Panels 1 and 2 turn each genome into a point in the simplex. Panel 3 is the criterion proposed here: rounding sends nearby profiles to the same grid cell, and equal cells give equal digests, so a genome joins its group without being compared with any other sequence. Panel 4 contrasts the two ways of dividing the data and panel 5 the comparison that follows, and the bordered box at the end states what the two benchmarks report.

On notation: *i* and *i*^*′*^ run over sequences, *j* and *j*^*′*^ over features, *c* and *c*^*′*^ over classes, *u* over near-duplicate groups and *t* over optimisation steps; vectors and matrices are bold and their entries italic, so **f**_*i*_ is a profile and *f*_*ij*_ one of its entries. Two pairs are worth separating in advance: Δ^*d−*1^ is always the probability simplex and Δ**W** a parameter step, and *δ* is always the rounding resolution introduced below, never a variation.

### 2.1. Materials

Whole-genome SARS-CoV-2 sequences were downloaded in FASTA format from NCBI Virus^1^ on 12 July 2026, for twelve Pango lineages: B.1.1.7 (Alpha), B.1.351 (Beta), P.1 (Gamma), B.1.617.2 (Delta), B.1.1.529 (Omicron), JN.1, LP.8.1, XDE, XDV, XDV.1, XFG and XFY, giving 342,568 sequences. A repository changes over time, so the collection is archived exactly as retrieved and is publicly available;^2^ the date and that copy together fix what was used.

All work was done in Python 3: Biopython for parsing, NumPy and pandas for arrays, scikit-learn 1.6.1 for splitting and scoring, and PyTorch 2.2 [14] for the networks and all optimisers.

### 2.2. Cleaning and class selection

Each sequence was converted to upper case and stripped of gaps, whitespace and line breaks, then kept only if it was at least 25,000 nucleotides long and had at most 5% ambiguous, non-ACGT characters. Exact duplicates within a lineage were removed by hashing the cleaned string and keeping the first copy. Of the 342,568 sequences loaded, 86,343 were rejected — 3 for length, 33,030 for ambiguity, 53,310 as exact duplicates — leaving 256,225, or 74.80%.

Five lineages were then dropped. Three recombinants held too few sequences to support a per-class score: XFG with 368, XFY with 117 and XDV.1 with 113. The other two, XDE and XDV, held eight sequences each, and within each one all eight carried identical accession identifiers and byte-identical *k*-mer vectors — a defect in the source collection rather than biology, and a class made of one repeated record cannot be scored.

Seven lineages remained: Alpha (209,712), Delta (18,485), JN.1 (14,428), Gamma (9,081), Omicron (1,841), Beta (1,143) and LP.8.1 (921), together 255,611 sequences or 99.76% of the cleaned collection, so the exclusions cost very little data. The largest class is 228:1 larger than the smallest; Subsection 2.8 describes how that imbalance was handled.

### 2.3. From genome to vector

Each genome must become a vector of fixed length, twice over: the classifier takes vectors as input, and the near-duplicate test compares vectors rather than sequences. We slide a window of length *k* along the genome, one position at a time, and count how often each word appears. To keep the counts in a fixed order, each word *w* = *w*_1_*w*_2_ · · · *w*_*k*_ is read as a number in base 4, using the digits A ↦ 0, C ↦ 1, G ↦2, T ↦ 3, written *v*(·):

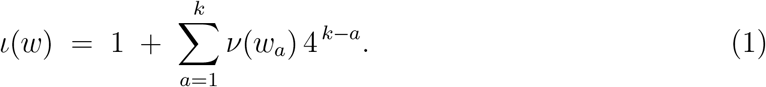

With *d* = 4^*k*^, the map *ι* of Eq. (1) is a bijection onto {1, …, *d*}, so each word has its own position and no two share one; feature indices run from 1 to *d* everywhere below. Windows holding an ambiguous character, such as N, are skipped. Write *W*_*k*_(*g*_*i*_) for the multiset of length-*k* windows of genome *g*_*i*_ that contain only A, C, G and T. Counting how many of them carry each index gives the count vector, whose *d* entries are

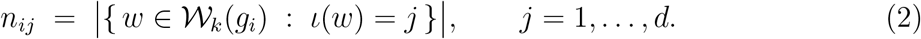

A longer genome gives larger counts everywhere, so the counts of Eq. (2) cannot be compared as they are. We divide by their total,

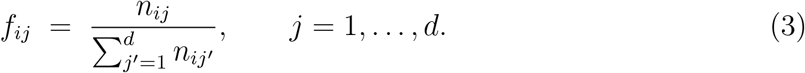

The divisor in Eq. (3) is |*W*_*k*_(*g*_*i*_)|, the number of valid windows rather than the genome length, because ambiguous windows were skipped. The results are non-negative and add to one,

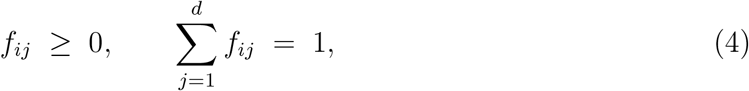

so each genome is one point **f**_*i*_ = (*f*_*i*1_, …, *f*_*id*_) in the probability simplex Δ^*d−*1^ ⊂ ℝ*d*. Equation (3) keeps only relative composition, and by Eq. (4) the *d* entries carry *d* − 1 free values.

The feature dimension is *d* = 4^*k*^: 64, 256 and 1024 for *k* = 3, 4 and 5. All results below use *k* = 4. Building the counts takes one pass over the sequence, so the cost does not depend on *d*.

### 2.4. Quantised profile hashing

Removing exact duplicates is not enough: many stored sequences differ at only a few positions and give almost the same vector under Eq. (3). The rule that decides when two genomes count as near duplicates must do more than answer yes or no for one pair, because the split keeps whole groups, not pairs. A distance test,

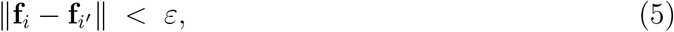

cannot do this. Closeness does not pass along a chain. Many small steps, each below *ε* in Eq. (5), add up to a large distance, so whether two sequences end up together would depend on the order in which pairs happen to be compared.

We propose *quantised profile hashing* (QPH), which rounds before it compares. Fix a resolution *δ* = 10^*−m*^ with *m* a positive integer, and round every entry of the frequency vector to the nearest multiple of *δ*, ties upward,

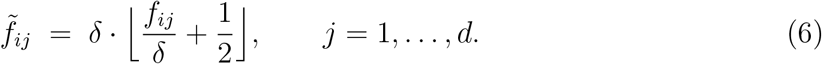

Write 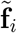 for the vector these entries form.

#### Definition 1 (Near duplicate)

Genomes *i* and *i*^*′*^ are *near duplicates at resolution δ*, written *i* ~_*δ*_ *i*^*′*^, when their rounded profiles agree in every coordinate,

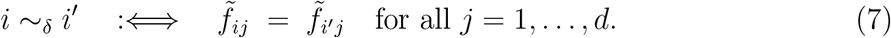

#### Proposition 1.

*The relation* ~_*δ*_ *of Eq*. (7) *is an equivalence relation. Its classes therefore partition the collection into disjoint groups G*_1_, …, *G*_*U*_, *and every genome lies in exactly one of them*.

#### Proof

By Eq. (6) every genome *i* has exactly one rounded profile 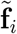, and Eq. (7) says *i* ~_*δ*_ *i*^*′*^ precisely when 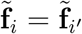. Equality of vectors in R*d* is reflexive, symmetric and transitive, and ~_*δ*_ is its pullback along 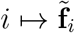, so it inherits all three properties. The classes of an equivalence relation are disjoint, non-empty and cover the collection. □

This is what QPH gains over the distance test. Each genome reaches its group through its own rounded vector, never through a comparison with a neighbour, and by Proposition 1 the groups cannot overlap — which is what the split requires and what the non-transitive Eq. (5) cannot deliver.

#### Remark 1

Rounding does not preserve Eq. (4): the entries of 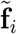 need not sum to one, so 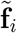 need not lie in Δ^*d−*1^. Nothing below uses more than equality of these vectors, so the departure is of no consequence.

In practice QPH hashes each rounded vector with BLAKE2b, and sequences sharing a digest go in the same group; a cryptographic hash is used because the interpreter’s built-in hash is randomised at every start and would not reproduce. We set *m* = 4, that is *δ* = 10^*−*4^, matched to the effect of a single substitution: it rewrites only the *k* windows covering the altered position and so moves at most 2*k* entries of Eq. (3), each by 1*/* |*W*_*k*_(*g*_*i*_)| ≈3 × 10^*−*5^ for a SARS-CoV-2 genome with |*W*_*k*_(*g*_*i*_)| ≈3 × 10^4^ valid windows, and by at most *k/* |*W*_*k*_(*g*_*i*_)| where the affected windows repeat a word. The typical shift is below *δ*, so genomes differing at one site round together unless an affected entry lies that close to a bin boundary, while genomes differing at many sites separate.

One caution: a smaller *δ* does not always split groups further. Entries 0.14999 and 0.15001 round apart at *δ* = 10^*−*1^ but together at *δ* = 10^*−*2^, while 0.10 and 0.11 do the opposite. Neither grouping refines the other, so *δ* must be fixed in advance and not tuned on results.

### 2.5. Splitting the data without leakage

Let (*T, V, E*) be a split of the sequences into pairwise disjoint training, validation and test parts. We measure contamination by the share of test sequences having a near duplicate in training,

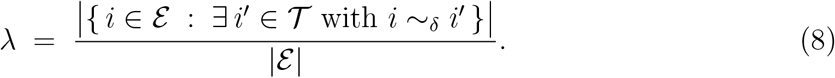

In Eq. (8), *λ* = 0 means no test sequence has a twin in training and *λ* = 1 means all of them do. Random splitting places sequences one by one, so nothing keeps *λ* small. We place groups instead. With *G*_1_, …, *G*_*U*_ the groups of Proposition 1, each group must sit inside one part only,

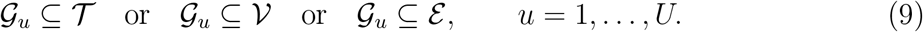

#### Proposition 2

*Any split obeying Eq*. (9) *has λ* = 0.

#### Proof

Suppose some *i* ∈*ε* had a near duplicate *i*^*′*^ ∈*T*. By Definition 1 and Proposition 1 they lie in the same group *G*_*u*_. Then *i* ∈*G*_*u*_ ∩ *ε*, so Eq. (9) places *G*_*u*_ inside *ε*; and *i*^*′*^ ∈ *G*_*u*_ ∩ *T*, so the same condition places *G*_*u*_ inside *T*. Then ∅ ≠*G*_*u*_ ⊆ *ε* ∩ *T*, contradicting disjointness. The numerator of Eq. (8) is therefore zero. □

The value of Proposition 2 is that *λ* = 0 follows from how the split is built, not from a measurement made afterwards. Its accuracy depends on Definition 1, whose two mistakes are not equally serious: putting unrelated sequences in one group only makes the split stricter and costs a few sequences, whereas separating two nearly identical vectors that fall on opposite sides of a rounding boundary is what lets contamination through. What we report is therefore a low estimate.

Groups are assigned to *K* folds so that every fold reproduces the class proportions of the whole collection as closely as the groups allow. Writing *N*_*uc*_ for the number of sequences of class *c* in group *G*_*u*_, and *ϕ* : {1, …, *U*} → {1, …, *K*} for the fold assignment, the map *ϕ* minimises

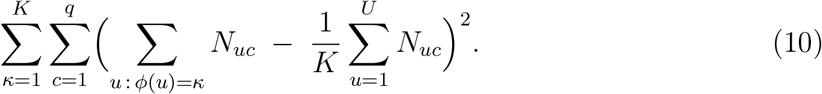

The inner sum of Eq. (10) counts class *c* in fold *κ*, and the term subtracted is what an even spread would give. Because *ϕ* acts on groups, Eq. (9) holds automatically.

Minimising Eq. (10) exactly is NP-hard: with one class and two folds it is the partition problem. We use the usual greedy assignment, taking groups in order of decreasing size and placing each in the fold furthest below target. This affects only class balance; it cannot break Eq. (9), so *λ* = 0 holds regardless. The scheme runs twice. The first pass uses *K* = 7 and takes one fold as the test set; the second uses *K* = 6 on the rest and takes one fold as the validation set. The group-aware split holds 182,585 training, 36,545 validation and 36,481 test sequences.

Running the same scheme with the grouping step omitted gives the random split used for comparison, at 182,579/36,516/36,516. It ignores the groups entirely, so nothing in it keeps *λ* small. Subsection 3.1 reports how large *λ* becomes, and Subsection 3.2 what each split then reports.

### 2.6. Feature normalisation

The networks below contain no normalisation layers, so input scaling is handled in the data. Two steps are applied. The first is Eq. (3), computed per sample. The second scales each feature,

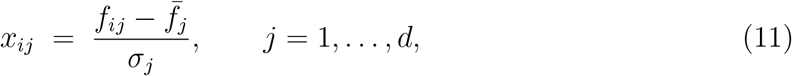

where the mean 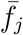 and the standard deviation *σ*_*j*_ in Eq. (11) come *from the training split alone* and are then applied unchanged to the other splits. Using the whole dataset would let held-out data influence the training representation. A feature with *σ*_*j*_ *<* 10^*−*8^ is left at *x*_*ij*_ = 0 instead of divided by a near-zero number.

Being a fixed coordinatewise affine map, Eq. (11) neither merges nor separates profiles; quantised profile hashing is applied to **f**_*i*_ before this step, so the grouping does not depend on statistics of the training split.

This step matters. Without it the entries of Eq. (3) are of order 1*/d*, so the input scale shrinks as *d* grows; under those conditions a two-layer version of this network never left the majority-class solution at any learning rate tried, whereas with Eq. (11) it passed a macro F1-score of 0.99 within one epoch.

### 2.7. The classifier

The classifier turns a standardised vector **x**_*i*_ ∈ ℝ*d* from Eq. (11) into *q* = 7 logits by one affine map,

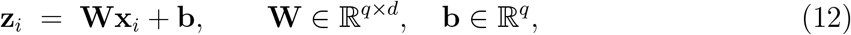

with *qd* + *q* parameters: 455 at *k* = 3, 1,799 at *k* = 4 and 7,175 at *k* = 5. Equation (12) is the entire model, kept simple on purpose because what is being tested is the split, not the architecture, and because a simple model makes the curvature exact rather than approximate, as Subsection 2.9 shows, so the second-order rules can be compared without that extra source of doubt.

The logit vector of sample *i* is **z**_*i*_ ∈ ℝ*q* with entries *z*_*ic*_; where the sample index is dropped it is **z** with entries *z*_*c*_. The softmax sends these logits to a probability over the *q* classes,

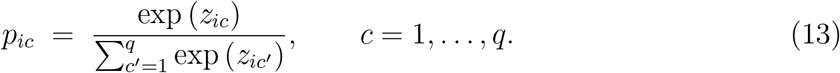

The entries of Eq. (13) are positive and sum to one over *c*, so *p*_*ic*_ is the probability the model assigns class *c* to sample *i*. Write *y*_*i*_ ∈{1, …, *q*} for the true class of sample *i* and 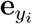 for its one-hot encoding,

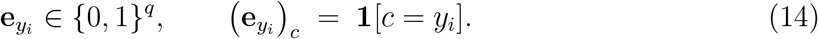

where **1**[] is the indicator, equal to 1 when its condition holds and 0 otherwise. The loss of sample *i* is the negative log probability of its true class; summing against the one-hot target and dropping the terms it sets to zero,

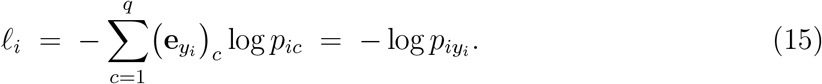

Equation (15) depends on **W** and **b** through **z**_*i*_, which is the route the derivatives of Subsection 2.9 follow. The network uses no batch normalisation, layer normalisation or dropout; the reason is a requirement of the second-order rules, given with Eq. (28), and Eq. (11) takes over the scaling role those layers would play.

### 2.8. Handling of class imbalance

Each sample’s loss is weighted by class, so that small classes are not ignored. Plain inverse-frequency weights, *β*_*c*_ = *N/*(*qN*_*c*_) for *N* training samples, *q* classes and *N*_*c*_ members of class *c*, carry the full 228:1 imbalance. In early runs that made training unstable. We damp it with a square root,

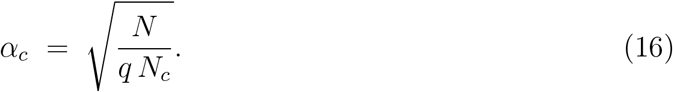

Equation (16) and the plain weights both give the largest weight to the smallest class. Writing *r* = max_*c*_ *N*_*c*_*/* min_*c*_ *N*_*c*_, their spans are

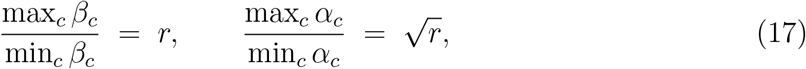

since a square root halves a ratio on a logarithmic scale. Here *r* = 228, so Eq. (17) gives 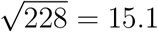 to one.

The two are related by 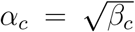, the geometric mean of the plain weight and the uniform weight 1. The plain weights carry exactly the mass of the unweighted training set; the damped ones carry at most as much,

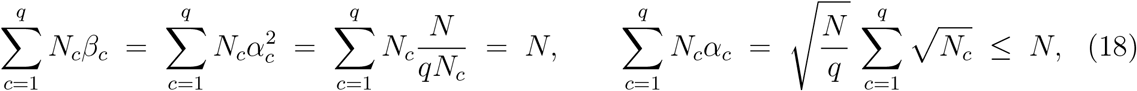

the inequality being Cauchy–Schwarz applied to 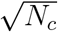 against 1, with equality exactly when all classes are of equal size. Damping therefore does two things in Eq. (18). It flattens the profile of weights across classes, which is the intended effect; and it multiplies the data term, but not the fixed *L*_2_ penalty below, by a constant under one. Since ζ, the learning rate and the weights are held at one value per rule across both partitions, that second effect is common to the two runs being compared and cannot account for any difference between them.

On a mini-batch B of size *B* the objective is

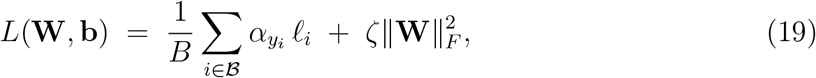

where *l*_*i*_ is the loss of Eq. (15) and the last term an *L*_2_ penalty with *ζ* = 10^*−*4^ on the weight matrix only, not on the bias.

### 2.9. Gradients and curvature of the classifier

Ten update rules are compared in Section 3: five standard first-order methods, named in Subsection 2.12, and five second-order rules that build their step from an estimate of the curvature of the objective. This subsection derives the quantities the latter need, for the single-layer network of Eq. (12). None of the ten is proposed as a contribution; all are instruments.

Start from one sample, dropping the sample index. Writing 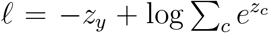 and differentiating with respect to a single logit gives

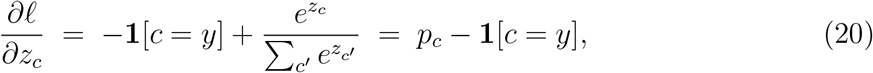

with **1**[·] the indicator of Eq. (14). In vector form Eq. (20) reads ∇_**z**_*l* = **p** − **e**_*y*_, the predicted distribution minus the one-hot target of Eq. (14). Differentiating Eq. (20) once more,

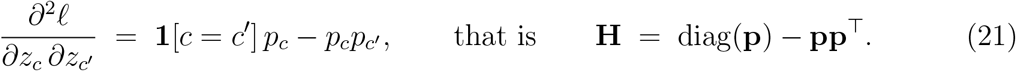

The matrix **H** in Eq. (21) is *q* × *q*, symmetric and positive semi-definite, and depends on the parameters only through **p**. Next, how the logits respond to the parameters: by Eq. (12), 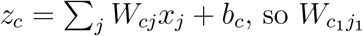 appears in *z*_*c*_ only when *c* = *c*_1_, and then with coefficient 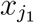,

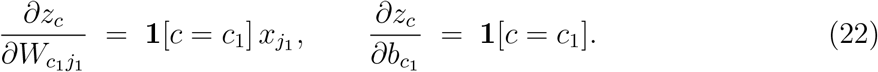

Equation (22) shows that every formula for the bias follows from the corresponding formula for the weights by setting 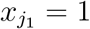.

Combining Eqs. (20) and (22) by the chain rule, and averaging over a mini-batch ℬ of size *B*, the gradient of the objective of Eq. (19) is

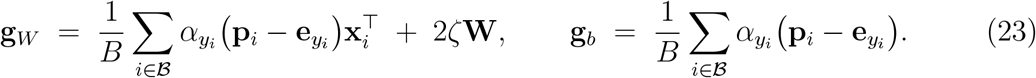

The curvature follows the same route. Writing **J** for the matrix of derivatives in Eq. (22), the per-sample Gauss–Newton matrix is **G** = **J**^*T*^**HJ**; substituting and indexing the parameters by the pair (*c, j*),

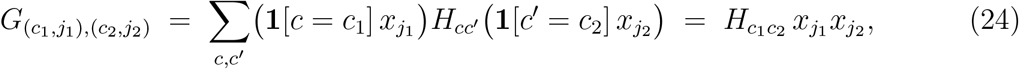

because the two indicators leave only the term *c* = *c*_1_, *c*^*′*^ = *c*_2_. Equation (24) says that the whole *qd* × *qd* curvature matrix is built from two small pieces,

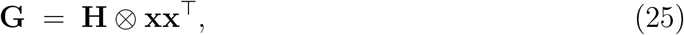

where ⊗ is the Kronecker product and vec stacks the *rows* of a matrix, so that *W*_*cj*_ sits at position (*c* − 1)*d* + *j* and the class factor comes first. That first factor is the *q* × *q* matrix of Eq. (21) and describes how the classes interact; the second is **xx**^*T*^, *d* × *d*, and describes how the features interact. Over a mini-batch the curvature of Eq. (19) is the class-weighted average 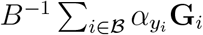 plus the 2*ζ***I** contributed by the *L*_2_ term.

One remark before the rules themselves. For a general network the Gauss–Newton matrix only approximates the Hessian [15], which carries an extra term in the second derivatives of the network output with respect to the parameters. Here **z** is affine in **W** and **b**, so those vanish and Eq. (25) is the exact Hessian of *l* with respect to **W**; with 2*ζ***I** it is the exact Hessian of the full objective.

### 2.10. The three diagonal rules

Forming and inverting **G** at every step (is imp)ractical: its weight block is *qd* × *qd*, that is 1792 × 1792 at *k* = 4, the solve costs *O* (*qd*)^3^, and the number of entries grows as *d*^2^; at *k* = 5 the block is already 7168 × 7168. The three rules DGN-GGN, DGN-EF and DGN-sqmean keep only its diagonal. They share one update engine and differ only in how that diagonal is estimated, so any difference between them is attributable to the estimate rather than to the step. The engine keeps a running average **v**_*t*_ of the estimate **s**_*t*_ supplied at step *t*, corrects it for its start at zero, and divides the gradient of Eq. (23) by it. Writing ***θ*** for either **W** or **b**, and **g**_*t*_ for the matching block of that gradient at step *t*,

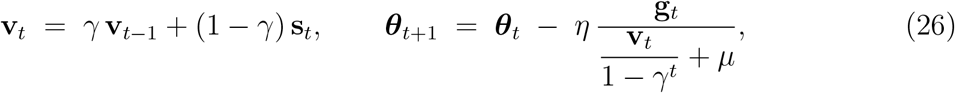

with decay *γ* = 0.95, learning rate *η*, damping *μ*, and all operations entrywise. The factor 1 − *γ*^*t*^ removes the downward bias that **v**_0_ = **0** would otherwise cause in the first steps, and gradients are clipped to a global norm of 1.0 before Eq. (26) is applied. The three paragraphs below give the entries of **s**_*t*_, with the step index dropped from their right-hand sides.

#### DGN-GGN

Take the diagonal of Eq. (25) directly. Setting *c*_1_ = *c*_2_ = *c* and *j*_1_ = *j*_2_ = *j* in Eq. (24), and noting from Eq. (21) that *H*_*cc*_ = *p*_*c*_(1 − *p*_*c*_), the estimate for the weight *W*_*cj*_ averaged over the batch is

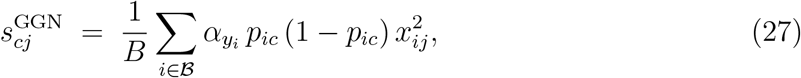

and the bias version follows by setting *x*_*ij*_ = 1. This is the most faithful of the three: Eq. (27) is the exact diagonal of the exact Hessian.

#### DGN-EF

Replace the Gauss–Newton matrix by the empirical Fisher, the average outer product of the per-sample gradients. By Eqs. (20) and (22) the per-sample gradient with respect to *W*_*cj*_ is 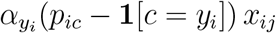, so squaring and averaging gives

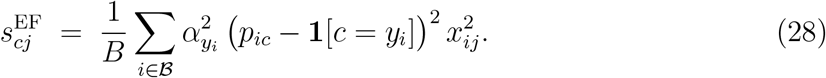

Equation (28) costs one matrix product, as Eq. (27) does, and needs no per-sample Jacobian, by the identity of Goodfellow [16]. That identity requires the loss to separate across samples, which is why the network carries no batch normalisation: batch statistics couple the samples and break the separation. The empirical Fisher is also not the Fisher information, and can differ from the curvature in ways that matter for optimisation [17]; comparing Eqs. (27) and (28) asks whether that difference shows up here.

#### DGN-sqmean

Use the square of the batch-mean gradient itself,

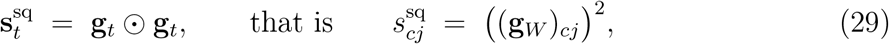

with ⊙ the entrywise product. Equation (29) is not the same as the average of the squared per-sample gradients used in Eq. (28). Write **g**_*i*_ for the gradient of the weighted per-sample loss 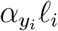, so that 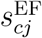 of Eq. (28) is the (*c, j*) entry of 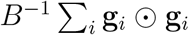 and 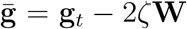, the batch gradient of Eq. (23) with the penalty term removed. Compared on this common data term the two differ by exactly the variance across the batch,

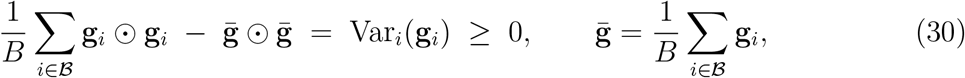

entrywise. The right-hand side of Eq. (30) is non-negative, so on that common term Eq. (29) can only underestimate Eq. (28), and by exactly the batch variance. The *L*_2_ penalty is added once to the batch objective rather than per sample, so it enters 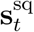 but has no per-sample counterpart in Eq. (28); at *ζ* = 10^*−*4^ its contribution is small beside the data term. Equation (29) is the quantity a common implementation of this family computes.

### 2.11. The two Kronecker-factored rules

The diagonal discards both off-diagonal factors of Eq. (25) at once. The rules KGN-diag and KGN-full keep the class factor whole, and differ from each other in what they do with the feature factor.

Averaging Eq. (25) over a batch gives a sum of Kronecker products, which is not itself a Kronecker product. Following the K-FAC approximation [18], we replace the average of the products by the product of the averages,

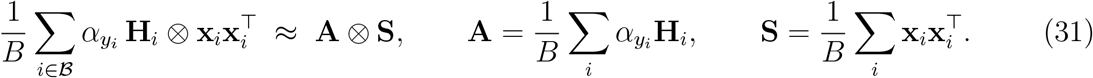

Both factors of Eq. (31) are averages of symmetric positive semi-definite matrices, so both stay invertible once damped. Substituting Eq. (21) gives the class factor in the form actually computed,

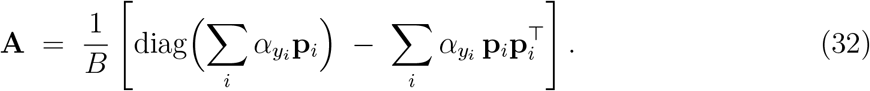

The class factor **A** of Eq. (32) is *q* × *q* with *q* = 7, so inverting it is negligible work.

The step solves the damped system (**A** ⊗ **S** + *μ***I**) vec(Δ**W**) = vec(**g**_*W*_). That system does not factor, so we use the standard factored damping, which splits *μ* evenly between the two factors:

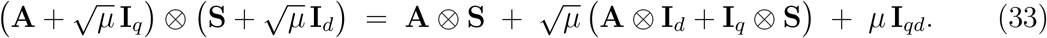

Equation (33) recovers *μ***I**_*qd*_ exactly and adds a cross term that vanishes with *μ*. For the row-major vec fixed at Eq. (25), the mixed-product identity reads (**P** ⊗ **Q**^*T*^) vec(**X**) = vec(**PXQ**) for conformable **P, Q** and **X**; applying it to the factored form turns the large solve into two small ones,

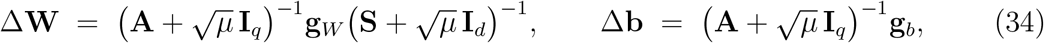

which is the rule KGN-full. By Eq. (22) the input of the bias is the constant 1, so its feature factor is the scalar *S*_*b*_ = 1 rather than a *d* × *d* matrix. We damp only the class factor for the bias, which leaves out the 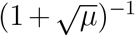 that factored damping would otherwise contribute and so scales the bias step by a fixed factor relative to the weight step; *μ* was selected with that convention in place.

The *d* × *d* inverse in Eq. (34) costs *O*(*d*^3^) peL.r step. KGN-diag keeps **A** exact but replaces **S** by its diagonal, whose entries are 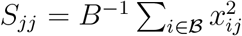, so that the right-hand solve becomes a division column by column,

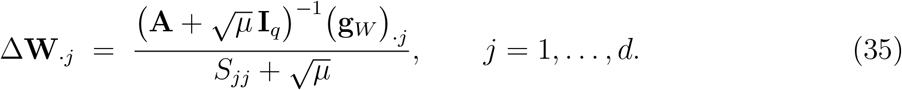

Both rules then take **W ← W −** *η* Δ**W** and **b ← b −** *η* Δ**b**. Comparing Eq. (35) with Eq. (34) isolates the feature factor, and comparing either with Eqs. (27) and (28) the class factor: the five second-order rules form a ladder in how much of Eq. (25) is retained, which is what makes them useful as instruments.

### 2.12. Baselines and verification

Five first-order optimisers serve as baselines: stochastic gradient descent with momentum 0.9, Adam [19], AdamW with decoupled weight decay 10^*−*4^ [20], RMSprop and Adagrad [21]. AdamW is the one rule that carries weight decay twice, since Eq. (19) already applies the *L*_2_ penalty to every rule and AdamW’s decoupled decay sits on top of it, as its definition requires; its configuration is identical on the two partitions, so the comparison drawn here is unaffected. With the five second-order rules of Subsections 2.10 and 2.11 this gives ten. All ten were written from their defining recursions and run inside one shared training loop, so batch order, initialisation, class weighting, clipping, the objective and the scoring are identical across methods; the update rule itself is the only thing that varies.

Each baseline was checked against the matching PyTorch routine on the same random positive-definite quadratic for 300 steps. Stochastic gradient descent, RMSprop and Adagrad agreed exactly; Adam and AdamW agreed to 2.9 × 10^*−*7^ relative, which is what differences in floating-point summation order produce. The library versions appear only in this check.

### 2.13. Training protocol

The batch size was 256 throughout, and hyperparameters were chosen in two stages. The first ran each optimiser once per configuration for 30 epochs at one fixed seed: five learning rates spanning five decades for each first-order method; six learning rates from 10^*−*6^ to 10^*−*1^ crossed with nine damping values from 10^*−*8^ to 1 for each diagonal method; and three learning rates crossed with three damping values for the two Kronecker rules. The second stage retrained each optimiser at its selected configuration for 60 epochs on three seeds (42, 123, 2024).

Model selection uses the validation split only: for every run the epoch with the highest validation macro F1-score is chosen and its parameters saved. The test split of 36,481 sequences takes no part in training or selection, is scored once from the saved parameters, and each saved file carries a fingerprint of the split so that it cannot be scored on a different partition unnoticed.

### 2.14. Evaluation metrics

The classes are very unequal in size, so plain accuracy is dominated by the largest and is reported only for comparison with earlier work. The main metric is the macro-averaged F1-score, built from the confusion matrix. Let 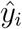 be the class predicted for item *i* in an evaluation set *D*, which is *V* during model selection and *E* for the reported scores, and let

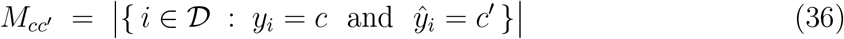

count the items of true class *c* predicted as class *c*^*′*^. From Eq. (36) the precision and recall of class *c* are

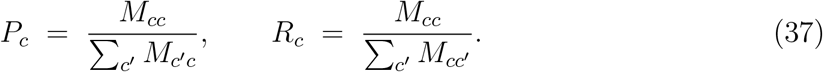

In Eq. (37) the denominator of *P*_*c*_ counts everything predicted as *c*, and that of *R*_*c*_ everything truly in *c*. We set *P*_*c*_ = 0 if nothing is predicted as *c*, and *R*_*c*_ = 0 if no item is in *c*; on the splits used here all seven classes are present in every evaluation set, so neither convention is ever invoked. The reported metric averages the per-class harmonic means,

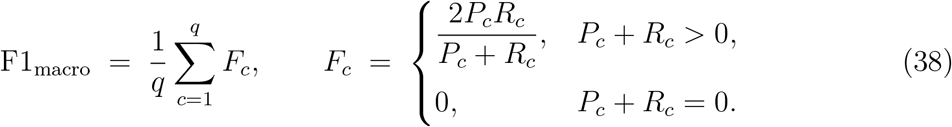

The second case of Eq. (38) avoids 0*/*0 for a class neither predicted nor present, and matches the limit. A harmonic mean lies between its arguments,

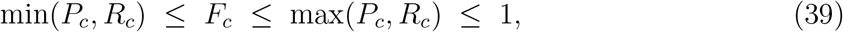

so by Eq. (39) a class scores well only when precision and recall are both high. Averaging with equal weights rather than by class size is what makes Eq. (38) sensitive to the small lineages: a failure on the 921 sequences of LP.8.1 is as visible as one on the 209,712 of Alpha. All results are means and standard deviations over the three seeds.

## 3. Results

All results use *k* = 4, that is *d* = 256, on the single-layer network.

### 3.1. How much contamination

Under ordinary stratified random splitting the leakage rate of Eq. (8) is *λ* = 0.0509. That is, 5.09% of test sequences had a near duplicate in the training set. The QPH-based split gives *λ* = 0, which by Proposition 2 follows from how the split is built rather than from a measurement made afterwards. Figure 2 shows the class distribution these numbers rest on, including the lineages that were excluded.

**Figure 2:**
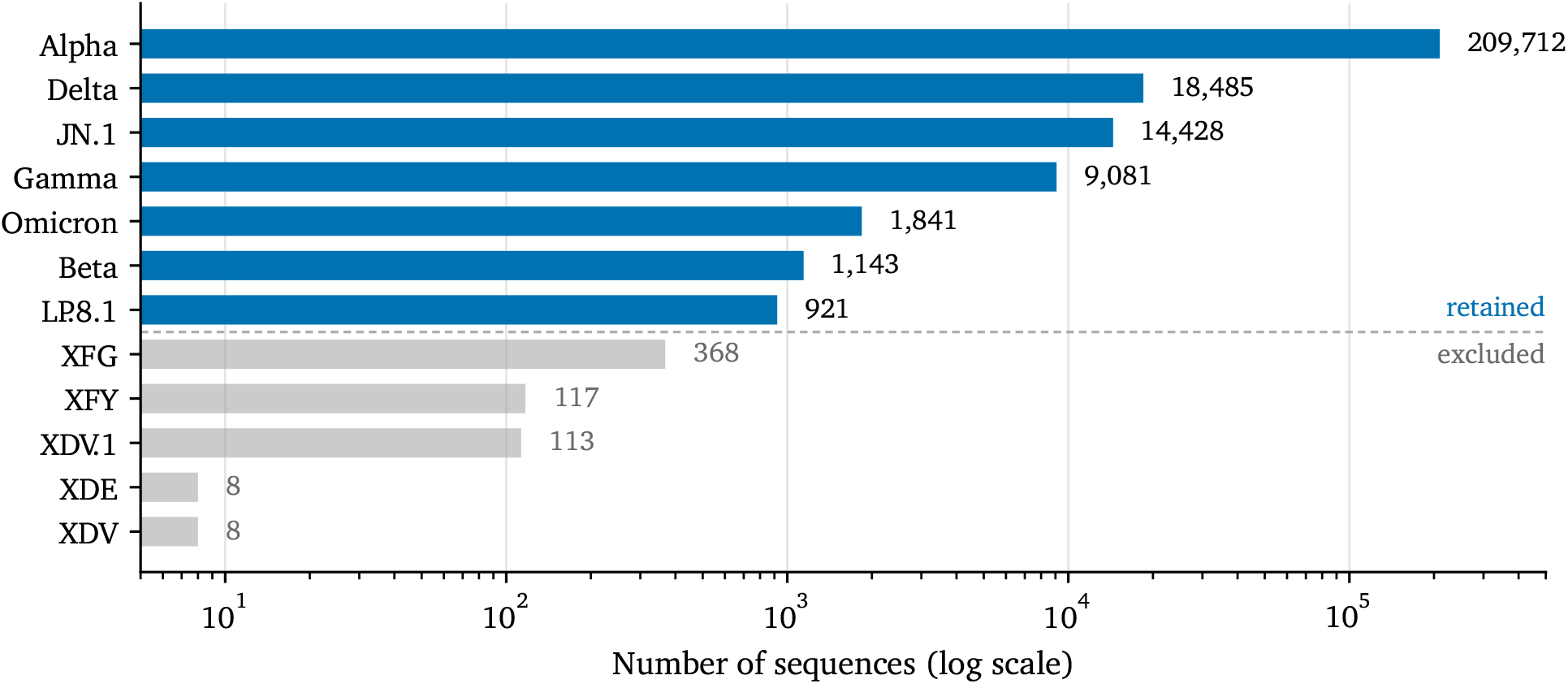
Sequence count of each lineage after cleaning, on a logarithmic axis. The seven retained lineages span the 228:1 imbalance between Alpha and LP.8.1; the five excluded ones are grey, below the dashed line — three too small to support a per-class score, two made entirely of repeated records

The number is small in absolute terms but not in context. Published methods in this field are separated by one or two percentage points [5, 6], so one test sequence in twenty is scored on material the model has already seen, on a benchmark whose verdicts turn on differences of that order. The two quantities are not the same kind of number: 5.09% is a share of the test set, while one or two points is a difference in score. Subsection 3.2 therefore measures the consequence directly instead of inferring it. Every result below comes from the group-aware split, so these numbers should not be compared directly with figures obtained elsewhere under random splitting..

### 3.2. What each benchmark reports

The ten rules were trained twice, once on each split, with hyperparameters, seeds, batch order, initialisation and epoch budget identical in the two runs; the partition is the only difference. Table 1 gives both outcomes, and three things follow.

**Table 1:** Held-out macro F1-score of ten update rules on the two splits, ordered by rank on the contaminated split. Mean and standard deviation over three seeds; each score is computed once, from the parameters saved at the epoch selected on the matching validation split. The *p* column is a two-sided Welch test against the leader of that column. ^*∗*^significant at 0.05 Separated from the leader at 0.05: 1 of 9 on the random split, 0 of 9 on the group-aware split. Ranks and rank correlations use full precision; scores are shown to five decimals, which is why three pairs of rules appear tied here.

| Update rule | Random split ($\lambda = 0.0509$) | | Group-aware split ($\lambda = 0$) | |
| --- | --- | --- | --- | --- |
| | Test macro F1 | $p$ | Test macro F1 | $p$ |
| DGN-EF | 0.99716 ± 0.00016 | — | 0.99642 ± 0.00018 | 0.711 |
| KGN-diag | 0.99716 ± 0.00031 | 0.991 | 0.99626 ± 0.00062 | 0.577 |
| KGN-full | 0.99700 ± 0.00071 | 0.730 | 0.99520 ± 0.00050 | 0.082 |
| RMSprop | 0.99692 ± 0.00030 | 0.319 | 0.99563 ± 0.00101 | 0.260 |
| Adam | 0.99657 ± 0.00043 | 0.127 | 0.99624 ± 0.00059 | 0.550 |
| Adagrad | 0.99642 ± 0.00077 | 0.236 | 0.99538 ± 0.00107 | 0.192 |
| DGN-sqmean | 0.99642 ± 0.00058 | 0.150 | 0.99566 ± 0.00022 | 0.182 |
| AdamW | 0.99636 ± 0.00048 | 0.092 | 0.99663 ± 0.00085 | — |
| DGN-GGN | 0.99598 ± 0.00116 | 0.218 | 0.99578 ± 0.00076 | 0.265 |
| SGD | 0.99587 ± 0.00036 | 0.014* | 0.99566 ± 0.00027 | 0.180 |

First, the contaminated benchmark produces a separation and the clean one does not. On the random split, stochastic gradient descent is significantly worse than the leader at the 0.05 level (*p* = 0.0143). On the group-aware split no rule is separated from the leader at that level, and the smallest *p*-value among the nine is 0.082. Contamination does not merely raise scores; it manufactures apparent significance.

Second, the ordering does not carry across. AdamW is eighth on the contaminated benchmark and first on the clean one, KGN-full moves the other way from third to last, and rules move by 3.2 positions on average. The two orderings are uncorrelated: Kendall *τ* = +0.022 (*p* = 1.00), Spearman *ρ* = +0.042 (*p* = 0.91). Figure 3 draws the movement. A ranking measured on the contaminated benchmark therefore carries no information about the ranking on the clean one.

**Figure 3:**
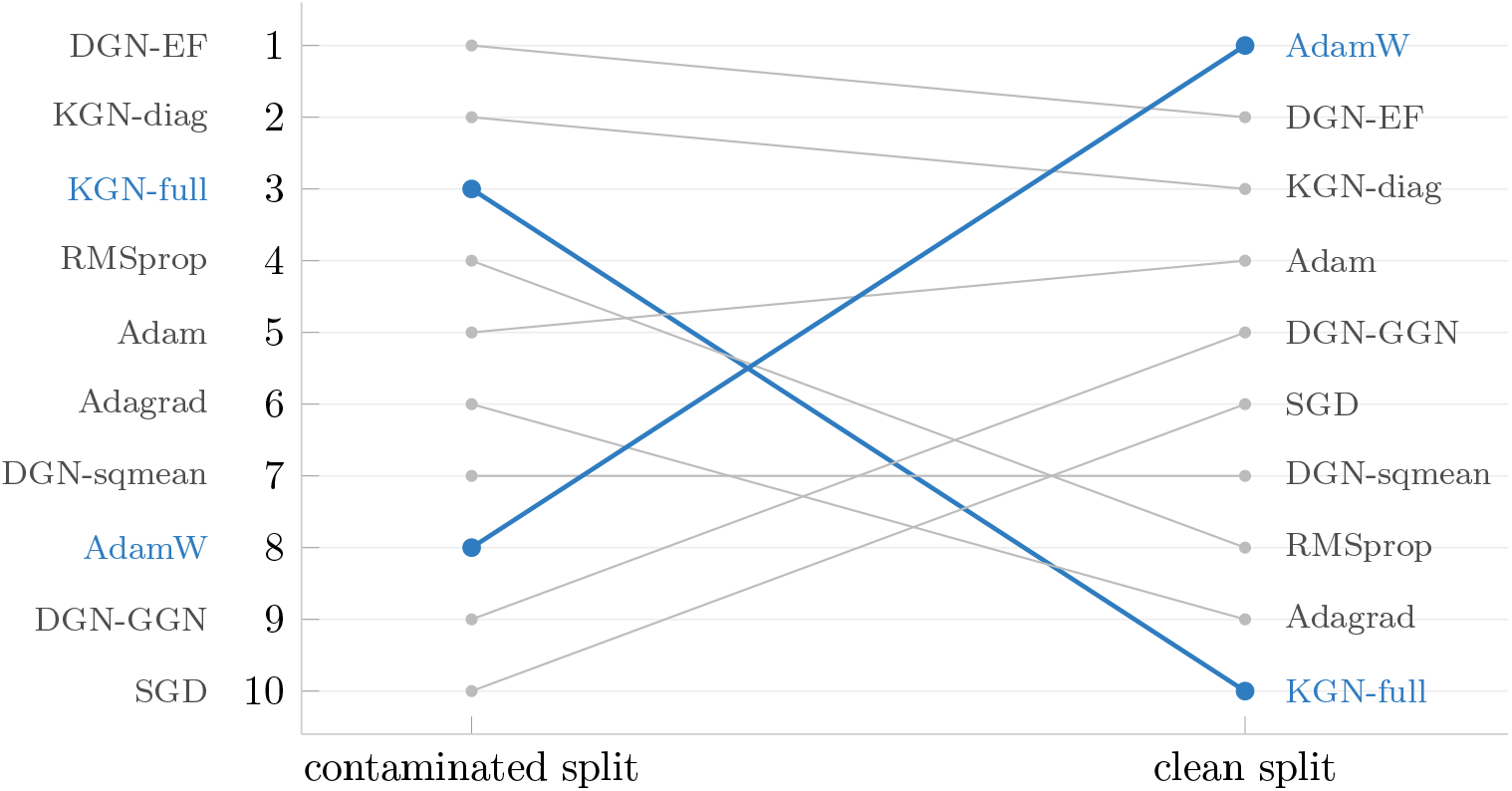
The ranking does not carry from a contaminated benchmark to a clean one. Each line follows one update rule from its held-out rank on the random split to its rank on the group-aware split, from Table 1, with everything but the partition held fixed. Rules move by 3.2 positions on average and the two orderings are uncorrelated, Kendall *τ* = +0.022; AdamW, in colour, rises from eighth to first and KGN-full falls from third to last.

Third, the contaminated split raises the score of nine rules of ten, by between 0.0002 and 0.0018 in macro F1; AdamW alone scores lower there, by 0.0003. The gain is therefore both small and unevenly distributed, and its spread is of the same size as the gaps between neighbouring rules, which is why the ordering is so easily rearranged.

Every rule exceeds a macro F1-score of 0.995 on both splits. By Eq. (25) the Gauss– Newton matrix is the exact Hessian for this model, so DGN-GGN is the only rule built from the true curvature; it ranks ninth on the contaminated split and fifth on the clean one, another way of seeing that neither position means much.

### 3.3. Per-class performance

Table 2 gives per-class held-out scores for the leading rule on the clean split. Since the macro average of Eq. (38) is driven by the smallest classes, those entries decide whether a split that never divides a group still leaves the rare lineages learnable. It does. Beta, Gamma and JN.1 are recovered without error on any seed; LP.8.1, the smallest class with 124 test sequences, reaches an F1-score of 0.9987 at perfect recall; and Alpha, which holds 82% of the data, reaches 0.9999. The weakest class is Omicron at 0.9803, also the only class whose scores move appreciably across seeds. Omicron and Delta have the closest *k*-mer profiles here and the errors concentrate between them, which is where a composition-based representation should fail first.

**Table 2:** Per-class held-out performance of AdamW, the leading rule on the group-aware split, at *k* = 4. Mean and standard deviation over three seeds.

| Lineage | Total | Test | Precision | Recall | F1-score |
| --- | --- | --- | --- | --- | --- |
| Alpha | 209,712 | 29,970 | $1.0000 \pm 0.0000$ | $0.9999 \pm 0.0000$ | $0.9999 \pm 0.0000$ |
| Delta | 18,485 | 2,612 | $0.9971 \pm 0.0012$ | $0.9981 \pm 0.0004$ | $0.9976 \pm 0.0004$ |
| JN.1 | 14,428 | 2,052 | $1.0000 \pm 0.0000$ | $1.0000 \pm 0.0000$ | $1.0000 \pm 0.0000$ |
| Gamma | 9,081 | 1,298 | $1.0000 \pm 0.0000$ | $1.0000 \pm 0.0000$ | $1.0000 \pm 0.0000$ |
| Omicron | 1,841 | 279 | $0.9809 \pm 0.0052$ | $0.9797 \pm 0.0110$ | $0.9803 \pm 0.0033$ |
| Beta | 1,143 | 146 | $1.0000 \pm 0.0000$ | $1.0000 \pm 0.0000$ | $1.0000 \pm 0.0000$ |
| LP.8.1 | 921 | 124 | $0.9973 \pm 0.0046$ | $1.0000 \pm 0.0000$ | $0.9987 \pm 0.0023$ |
| Macro average | | | $0.99663 \pm 0.00085$ | | |

That the rarest lineages are recovered as well as the most abundant one is the intended effect of the square-root damped weighting in Eq. (16). Overall accuracy for this model is 0.9997. The three smallest classes hold 549 of the 36,481 test sequences between them, so missing every one of them would still leave accuracy at 0.985. A metric that survives such a failure at three digits is not the metric to report here, which is why Eq. (38) is used throughout.

## 4. Discussion

### 4.1. What the ranking is worth

The two benchmarks disagree about which rule is best, and they disagree about whether any rule is best at all. Under random splitting 5.09% of test sequences carry a near duplicate from training. The scores this inflates move by at most 0.0018 in macro F1, an order of magnitude below the one-to two-point margins used to rank methods in this literature, and that is the point: an effect far too small to notice in a reported number is still enough to move rules by 3.2 positions and to leave the two orderings uncorrelated.

Contamination raises nine of the ten scores and lowers one, by amounts that differ from rule to rule and with a spread comparable to the gaps between neighbouring rules. It also produces a separation that the clean benchmark does not support. A reader of the contaminated table would conclude that stochastic gradient descent is significantly worse than the leader; the clean table gives no ground for that conclusion.

The useful quantity is therefore not a position in a table but *λ*, which costs one pass over the feature matrix and tells a reader how much of a reported score could have come from sequences the model had already seen.

### 4.2. Comparison with previous work

Table 3 lists reported results for sequence-based SARS-CoV-2 classification. Studies were found through Scopus and Europe PMC, and every figure was taken from the full text rather than the abstract.

**Table 3:** Reported results for sequence-based SARS-CoV-2 classification. acc.: accuracy. Where a study reports several experiments, the one closest to lineage or clade assignment is given; class counts refer to Pango lineages, GISAID clades or taxonomic ranks, depending on what is classified. The present study is deliberately absent, since placing one of its scores here would invite exactly the comparison this paper argues against.

| Study | Representation | Classes | Sequences | Reported |
| --- | --- | --- | --- | --- |
| Elsherbini et al. [22] | di/tri-nucleotide | 11 | 1,131,185 | 0.8788 acc. |
| Ullah et al. [23] | temporal conv. network | 8 | n.s. | 0.8836 acc. |
| Câmara et al. [10] | 1D CNN, raw sequence | 4 | 14,684 | 0.9194 acc. |
| Ávila Cartes et al. [7] | FCGR + CNN | 11 | 191,456 | 0.9622 acc. |
| Coutinho et al. [9] | sparse autoencoder | 8 | 1,896 | 0.9630 acc. |
| van Zyl et al. [5] | six AF methods + RF | 3,502 | 297,186 | 0.9780 acc. |
| Yu et al. [6] | $k$-mer NV + GAT | 103 | 182,851 | 0.9636 F1 |
| Jamhuri et al. [24] | seq2int, spike protein | 6 | 24,000 | 0.9789 F1 |
| Awe et al. [25] | CNN-BiLSTM, spike gene | 5 | 35,800 | 0.9991 acc. |

Three differences stand out, none of them a claim about which method is stronger. First, the class sets are not comparable: these studies classify anything from taxonomic ranks to 3,502 Pango lineages, with scores from 0.879 to 0.999, and two of the nine use the spike protein alone [25, 24]. An accuracy over 3,502 classes and one over seven measure different tasks. Second, only one of the nine reports a macro-averaged F1-score [6]; the rest report accuracy, or an F1-score whose averaging is unstated. Here the largest class holds 82% of the sequences, so a classifier that answered Alpha every time would score 0.82 on accuracy and 0.13 on macro F1, and accuracy alone cannot show whether the rare lineages were found. Third, seven of the nine state how they split their data, and the procedures are sound in the usual sense — stratified sampling, hold-out sets, cross-validation, and in Yu et al. [6] a split of each lineage by date. What none of them does is group near-duplicate sequences before splitting; stratification balances classes, it does not separate near-identical genomes.

One entry shows why this matters even when the split is stated. Awe et al. [25] use stratified sampling and report a test accuracy of 99.91 ± 0.03% against a training accuracy of 99.74 ± 0.11%. A test score above the training score can come from strong regularisation; it is also what happens when the test set holds sequences close to the training set, and reporting *λ* would settle the question directly.

### 4.3. Limitations

Four limits apply. The evaluation covers one collection of SARS-CoV-2 genomes and seven lineages, so rare and newly emerging recombinant lineages are not covered.

QPH works on composition rather than alignment, and can err in two directions. It may group genomes that are not close, either because their compositions happen to agree or because their differences leave composition unchanged; that error only makes the split stricter and costs a few sequences. It may also fail to group two nearly identical profiles that fall on opposite sides of a rounding boundary, and that is the error which lets contamination through, so what we report is a lower estimate.

Hyperparameters were selected once, on the group-aware split, and reused unchanged on the random split; retuning on each would confound the partition with the tuning.

Finally, *δ* and the choice *k* = 4 fix which sequences count as near duplicates. The same partition carries a higher leakage rate measured at *k* = 3 and a lower one at *k* = 5, because a longer word gives a finer profile, so the criterion must be applied at the representation the model actually receives.

## 5. Conclusions

Alignment-free lineage assignment is almost always evaluated with stratified random splitting, and methods are ranked by margins of one or two percentage points. On 255,611 SARS-CoV-2 genomes from seven lineages, 5.09% of test sequences carry a near duplicate from the training set under that protocol, so one test sequence in twenty is scored on material the model has already seen.

The remedy is cheap. Quantised profile hashing finds near duplicates in *k*-mer feature space by rounding each frequency vector and hashing it, in a single pass and without comparing any sequence with another. Rounding is what makes the groups well defined, a distance threshold being non-transitive and yielding no partition, and using those groups as units that are never split removes the contamination by construction.

Removing it changes what the benchmark reports. Ten update rules were trained twice, identically except for the partition. The contaminated benchmark separates one rule from the leader at the 0.05 level and the clean benchmark separates none, while the two orderings are uncorrelated and the leader of one benchmark ranks eighth on the other. A ranking obtained under contamination therefore says nothing about the ranking without it. The quantity worth reporting beside a score is *λ*, the leakage rate of the split.

Two directions follow. The first is to apply QPH to benchmarks already published, which needs only their feature matrices and splits. The second is to extend the criterion beyond composition, using alignment or sequence sketches to bound the error that remains.

## Acknowledgments

The authors thank the depositors of SARS-CoV-2 genome sequences to NCBI Virus, without whose submissions this study would not have been possible.

## Data availability

The sequence collection was retrieved from NCBI Virus on 12 July 2026 and is archived unchanged in a public repository, available at https://www.kaggle.com/datasets/edumath/sars-cov-2-variant-july-2026. The code, both cached partitions, and the sixty trained models are deposited in Zenodo at https://doi.org/10.5281/zenodo.21912227, so that every result can be reproduced without repeating any training. That identifier resolves to the latest version; the release used here is https://doi.org/10.5281/zenodo.21912228.

## Declaration of generative AI use

During the preparation of this work the authors used Claude to help implement and verify the training code and edit the language of the manuscript. The study design, the choice of methods, the interpretation of the results, and every claim made here are the authors’ own. All AI-assisted output was checked against the recorded data and the source code before use. The authors reviewed, edited the content and take full responsibility for it.

## CRediT Authorship Contribution Statement

**Mohammad Jamhuri:** Conceptualization, Methodology, Software, Formal Analysis, Investigation, Data Curation, Visualization, Writing – Original Draft, Writing – Review & Editing, Funding Acquisition. **Andy Irawan:** Validation, Formal Analysis, Investigation, Writing – Review & Editing, Supervision.

All authors have read and agreed to the published version of the manuscript.

## Footnotes

1 https://www.ncbi.nlm.nih.gov/labs/virus/vssi/

2 https://www.kaggle.com/datasets/edumath/sars-cov-2-variant-july-2026

